# Gut microbiota alterations due to fecal transplant

**DOI:** 10.1101/2020.07.31.231233

**Authors:** Evgenii I. Olekhnovich, Artem B. Ivanov, Vladimir I. Ulyantsev, Elena N. Ilina

## Abstract

**Background:** Fecal microbiota transplantation (FMT) is currently used to treat recurrent clostridial colitis and other diseases. However, neither the therapeutic mechanism of the FMT nor the mechanism that allows the donor bacteria to colonize the intestine of the recipient has yet been described. Moreover, FMT is a great model for studying the ecology of host-associated microbial communities. This creates the need for experimentation with approaches to metagenomic data analysis which may be useful to the interpretation of observed biological phenomena.

**Methods:** Here the RECAST (Recipient intestinE Colonisation AnalysiS Tool) computational approach is presented, which is based on the shotgun reads sorting process in accordance with their origin in recipient metagenome. Using the RECAST algorithm, taxonomic/functional annotation, and machine learning, the shotgun metagenomic data from three FMT studies including healthy volunteers, patients with clostridial colitis and metabolic syndrome were analyzed.

**Results:** According to the analysis results, the colonizing and remaining microbial diversity in the post-FMT recipient metagenomic samples is clearly separated from the non-colonizers and lost. It is well explained by higher relative abundance in donor/pre-FMT recipient, Human Microbiome project metagenomes, and taxonomy. Moreover, the colonizing and remaining microbes are associated with lantibiotic and tetracyclines resistance genes.

**Conclusion:** Based on obtained results, the previously proposed “core” human gut microbiome concept may be elaborated. The top microbes of gut microbiota form “cores”, which, moreover, are mutually integrable between humans. Also, we assume that redistribution of microbial diversity in post-FMT recipients’ metagenomes is due to competition of donor/recipient microbes and to host immunity. The associations of top gut microbes with lantibiotic/antibiotic resistance can be related to gut microbiota colonization resistance phenomena or anthropogenic impact.

## Background

Gut microbiota is a large community of microorganisms and viruses, which is a key player in the host body metabolism. Metabolic functions of the gut microbial consortia are associated with support of physiological homeostasis, synthesis of vitamins and amino acids, short-chain fatty acids and other essential functions [Belkaid and Harrison, 2017]. Development of gut microbiota starts from the moment of birth and may depend on some important events, a few of which can be distinguished: the way of birth (vaginal or cesarean section), maternal microbiota transmission, feeding (breastfeeding or artificial) [Bäckhed et al., 2015; Ferretti et al., 2018; Shao et al., 2019], early antibiotic therapy [Korpela et al., 2016] et al. Moreover, bacteria could enter the intestines from the environment with food [Milani et al., 2019] and drinking water [Hansen et al., 2018]. These factors form a microbial community that may contain both common and unique members for different people. Moreover, the community could be changed over time.

Fecal microbiota transplantation (FMT) is currently used to treat recurrent Clostridioides difficile infection (CDI) and other diseases. A number of studies are ongoing in inflammatory bowel diseases (IBD) (Crohn’s disease, nonspecific ulcerative colitis) and metabolic disorders. The changes of the intestinal microbiota during FMT by colonization with donor bacteria are described in the case of IBD [Li et al., 2016; Kumar et al., 2017; Lee et al., 2017; Smillie et al., 2018] and healthy volunteers [Goloshchapov et al., 2019]. However, neither the therapeutic mechanism of the FMT nor the mechanism that allows the donor bacteria to colonize the intestine of the recipient has yet been described. This creates the need for experimentation with approaches to metagenomic data analysis. In several studies donor bacteria colonizations are described based on various strategies such as single-nucleotide variants (SNV) detection [Li et al., 2016; Smillie et al., 2018; Kumar et al., 2017] and genome-resolved metagenomics (GRM) [Lee et al., 2017].

Here we present an alternative technique which allows the study of donor/recipient microbial reshaping due to FMT. This approach is based on the separation of donor’s and recipient’s metagenomic reads and allows creation of reads “baskets” by origin: came from donor, came from recipient before transplantation, origin is unknown etc. We used taxonomic annotation of reads within the certain “basket” for study the behavior of donor/recipient bacteria alteration during fecal transplant. Based on functional annotation, we predicted some features which can be associated with colonization. The obtained results confirm and supplement the previous findings on similar topics.

## Methods

### Reads classification algorithm

We developed the **RECAST**(**R**ecipient intestin**E C**olonisation **A**nalysi**S T**ool) algorithm based on MetaCherchant source code [Olekhnovich et al., 2018] to compare two metagenomes and distinguish which reads of one metagenome are found in another. It takes as input two samples with paired-end reads in fasta or fastq format. One of the samples is referred to as queried, another as analysed. On the first stage, the program retrieves all k-mers from the queried metagenome and saves the quantity of each k-mer in a data structure referred later in this article as de Bruijn graph. On the second stage, each pair of reads from analysed metagenome is searched against the queried metagenome. Each read is split into k-mers, which are searched for in the de Bruijn graph. As a result, mean depth coverage of a read by k-mers as well as breadth coverage of a read is obtained. Latter is defined as a proportion of positions in read, covered by k-mers from de Bruijn graph. Then theoretical estimation of breadth is used to classify each read as found or not found in the de Bruijn graph.

Given the mean depth coverage, the theoretical breadth coverage estimation is required for comparison to the calculated one. Let us assume that the amount of k-mers covering a fixed position in the read obeys Poisson distribution with probability mass function

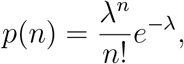

where *λ* equals mean depth coverage (*λ* = meanCoverage). This assumption is reasonable considering the fact that read is covered evenly and there are no jumps in coverage. Hence, the probability of position in read being not covered equals

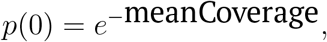

which equals the proportion of not covered positions in read for the long enough one. Consequently, theoretical breadth can be found as

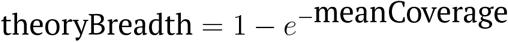

[Bankevich and Pevzner, 2018].

Having calculated this, we are next to define the confidence interval, which will contain the reads classified as found. It is approximated based on the central limit theorem. Breadth coverage is in the range (theoryBreadth − *δ*; theoryBreadth + *δ*) at 95% confidence level for

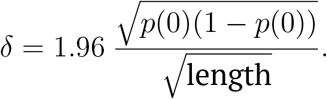

Also we introduce two user-defined thresholds for read breadth coverage. If breadth ≥ high_threshold (0.9 by default) read is supposed to be well-covered in the metagenome, otherwise if breadth *<* low_threshold (0.4 by default) read is supposed to be poorly covered in the metagenome. For more robust classification, two de Bruijn graphs with different lengths of k-mers are built (31 and 61 were used for analysis). Read is classified as **found** if for both values of k it satisfies high threshold AND falls within the confidence interval. If read is not **found** in any of two graphs and does not satisfy low threshold for at least one graph it is classified as **not found**. All other reads are hard to analyze due to being controversially covered, thus they are not used in subsequent analysis. During the processing of paired-end reads, reads from some pairs can be classified into different categories. This indicates the discrepancy between the classification and paired nature that might have been caused by small genome variations or sequencing errors. These read pairs are not credible and are excluded from further analysis. The schematic workflow of the algorithm is presented in Figure 1A.

**Figure 1.**
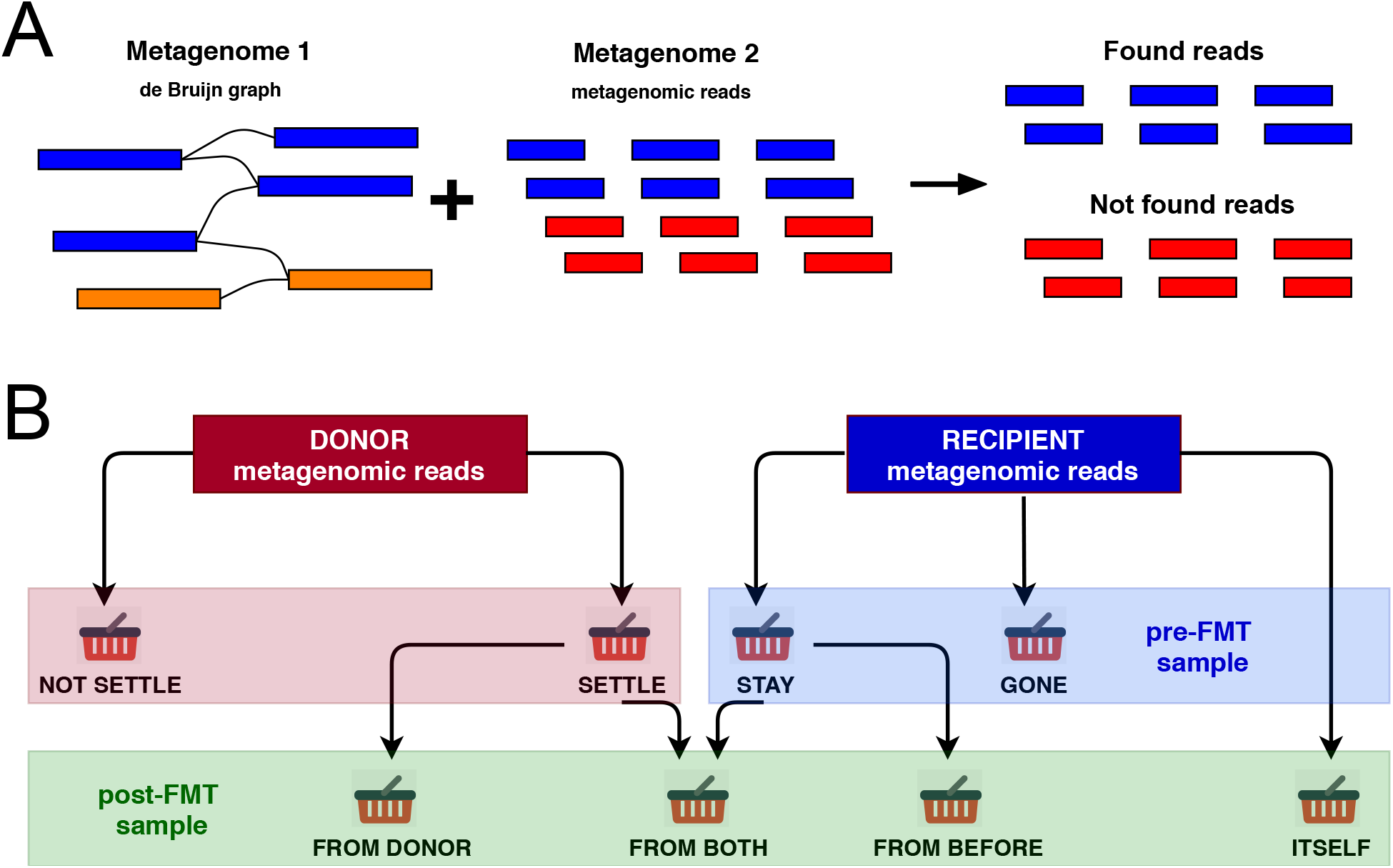
The reads classification algorithm workflow. **(A)** The basic scheme of reads classification algorithm. **(B)** The scheme of obtaining metagenomic reads “baskets”.

### FMT reads classification by origin scheme

A design of FMT experiments to study the behavior of the gut microbiota usually involves the collection and subsequent stool samples sequencing of donor, recipient before FMT (pre-FMT sample), and recipient after FMT (post-FMT sample). The RECAST algorithm takes as input every two out of three described above metagenomic samples and split reads from each metagenome into different categories based on the origin in the recipient’s metagenome.

Firstly, donor reads are queried against post-FMT recipient sample to generate categories **settle** and **not settle**. Secondly, pre-FMT recipient reads are queried against post-FMT recipient sample to generate categories **stay** and **gone**. Thirdly, post-FMT recipient reads are queried against donor sample and split into two temporary categories: found and not found. Finally, each category is queried against pre-FMT sample to split post-FMT reads into four categories: came **from both** – reads found in both donor and pre-FMT samples, came **from donor** – reads found only in donor sample, came **from baseline** – reads found only in pre-FMT sample and came **itself** – reads not found in donor nor in pre-FMT samples. The scheme of the produced reads categories is presented in Figure 1B.

### Simulated and real metagenomic data

#### Simulated data

We used simulated metagenomic time series created from bacterial genomes (see Supplementary Table S1) to assess the quality of reads classifier. 100 bp paired-end reads were sampled using MetaSim [Richter et al., 2008] with user-defined error model (see Supplementary Figure S1A). For additional “strain” simulations *Lactobacillus johnsonii* NCC 533 and *L. reuteri* DSM 20016 bacterial genomes available in the NCBI database were used. The above strains were grouped by nucleotide proximity using Mash distance [Ondov et al., 2016].

#### Real metagenomic data upload and quality control

The experimental FMT data used in this study are the longitudinally collected recipient metagenomes (one time point before transplantation and several after), as well as associated donor metagenomes. All available at the start of the study whole-genome sequencing (WGS) metagenomes containing both donor and recipient samples were selected. An additional criterion for data selection was the presence of aligned sampling points between recipients.

The data from FMT-allogeneic experiments in healthy volunteers [Goloshchapov et al., 2019] (healthy FMT), patients with Clostridioides difficile infection [Lee et al., 2017] (CDI) and metabolic syndrome [Vrieze et al., 2012] (MS allogeneic) were used. Additionally, autologous FMT data were used on patients with metabolic syndrome [Vrieze et al., 2012] (MS autologous) and from healthy people without interventions [Voigt et al., 2015] (Control) as comparison groups. Description of datasets is presented in Table 1 and Supplementary Table S2. For additional computational experiments 139 metagenomic stool samples from HMP 2012 dataset [Pasolli et al., 2016] were used. It is worth noting that the V1 volunteer (healthy FMT dataset) received two FMTs from one donor. The second FMT was carried out 38 days after the first FMT [Goloshchapov et al., 2019]. In total, 249 metagenomic stool samples were used in the study.

**Table 1.** Metagenomic datasets used in the study.

| Dataset | No. all individuals/<br>samples | No. donors/<br>samples | No. recipients/<br>samples | No. reads per<br>metagenome<br>(mean ± SD),<br>mln | Sequencing<br>platform<br>(read length,<br>bp) |
| --- | --- | --- | --- | --- | --- |
| Healthy | 4/17 | 1/3 | 3/14 | 23.3 ± 3.7 | Illumina (250) |
| CDI | 3/10 | 1/4 | 2/6 | 57.2 ± 13.4 | Illumina (150) |
| MS<br>allogeneic | 8/30 | 3/5 | 5/25 | 56.6 ± 18.0 | Illumina (100) |
| MS<br>autologous | 5/25 | — | — | 53.1 ± 19.5 | Illumina (100) |
| Control | 5/27 | — | — | 73.1 ± 38.8 | Illumina (100) |

Raw metagenomic data were downloaded from public repositories using fastq-dump in the SRA Toolkit [Sherry and Xiao, 2012], quality assessment was performed with FastQC https://github.com/s-andrews/FastQC and technical sequences and low-quality bases (*Q <* 30) were trimmed with Trimmomatic tool [Bolger et al., 2014]. The human sequences from metagenomic samples were removed by bbmap [Bushnell, 2014]. Described reads preprocessing computational steps were implemented in Assnake metagenomics pipeline https://github.com/ASSNAKE. The preprocessing statistic results are presented in Supplementary Table S3.

After reads quality control, sorting using the RECAST algorithm by the categories as described above was performed. As a control, sorting was performed in the Control group. Each baseline metagenome was selected as a “donor sample” (while remaining metagenomes from this subject were not used in the analysis), as in the case of real clinical data. In total, the 10 sorts series were performed in real-FMT datasets and 5 in Control.

### Data analysis and visualisation

Metagenomic taxonomic profiling was performed using MetaPhlAn2 [Segata et al., 2012; Truong et al., 2015]. Additionally, for tracking donor-derived bacteria in recipient metagenomes metaSNV [Costea et al., 2017] profiling based on mOTUs2 pipeline database [Milanese et al., 2019] was used. Manhattan distance was applied as a dissimilarity metric, which was averaged over all detected taxa in the sample. The Random Forest algorithm was used to range metadata/taxonomic features contribution to colonization. For model building and associated data processing pandas, numpy, scikit-learn libraries for python 3 and jupyter-lab were used. For functional analysis HUMAnN2 [Franzosa et al., 2018] and KEGG database (release 2018-03-26) [Kanehisa et al., 2017] was used. To determine differences in functional profiles Songbird tool [Morton et al., 2019] implementation via Qiime 2 framework [Bolyen et al., 2019] was used. The effect size threshold was set to 2.

Additional statistical testing and visualization were performed using Wilcoxon signed-rank test, vegan package [Oksanen et al., 2013] (Bray-Curtis dissimilarity and metaMDS function with default parameters), and ggplot2 library https://ggplot2.tidyverse.org implemented for GNU/R.

## Results

### Simulation metagenomic datasets

Firstly, we generated several datasets with different sample size parameter (number of reads per metagenome = 500000, 1000000, 2000000) and run the classifier on them varying k-parameter (k = 31, 41, 51). Each donor, pre-FMT recipient and post-FMT recipient metagenomes consisted of 8 randomly sampled genomes with relative abundance proportional to the genome’s length. Further, the results of classification were assessed in terms of recall, i.e. the fraction of correctly classified reads among the total number of reads expected in each category, precision, i.e. the fraction of correctly classified reads among the total number of reads classified in each category, and F1-score, which is the harmonic mean of two above (see Supplementary Figure S1B). We can conclude that the coverage of species in metagenome sample has a significant influence on the classification results, which requires careful experiment design. At the same time, the choice of k-parameter changes the classification results only slightly, unlike the other bioinformatics applications (e.g. assembly), which simplifies the step of launching the program for the end user.

Secondly, we varied the relative abundance of species in sampled metagenomes to study the effect on classification. Relative abundance was drawn from exponential distribution and generated values for all datasets are shown in Supplementary Figure S1C. We generated several datasets with different sample size parameter (number of reads per metagenome = 500000, 1000000, 2000000, 4000000) and run the classifier on them varying k-parameter (k = 31, 41, 51). Consistent with the first experiment, coverage has a great impact on classification results, whether k parameter is less important (see Supplementary Figure S1D). As expected, more depth coverage was needed to obtain the same classification quality as in the first experiment, due to the fact that species with small relative abundance have less coverage than mean coverage per dataset. Thirdly, metagenomes consisting of multiple strains of one species with different intraspecies distances were examined to assess the quality of the classifier. Each metagenome consisted of 4 randomly sampled strain genomes with uniform relative abundance with varied numbers of reads per metagenome (200000, 500000, 1000000, 2000000). Datasets were classified with varied k-parameter (31, 41, 51), which had a little effect. The quality of classification is highly dependent on relative distance between strains. While distant strains (mash distance 0.020 − 0.038%) can be detected and separated with medium accuracy provided sufficient depth coverage, close strains (mash distance 0.012−0.025%) can be hardly distinguished. (Our error model implies the probability error of 0.02%, thus we can not tell any difference between an error and a close strain). The results are presented in Supplementary Figure S1E.

### Real metagenomic datasets

#### Description of FMT data using metagenomic analysis

The common mutually complementary techniques for basic analysis of real metagenomic data includes taxonomic annotation by MetaPhlAn2 [Truong et al., 2015] and microbial strains profiling by metaSNV [Costea et al., 2017]. Five longitudinal metagenomic datasets were analysed. In addition to the real FMT allogeneic data (HEALTHY, CDI and MS Allogeneic sample sets), two data sets were used as controls: longitudinal autologous FMT (MS Autologous) and longitudinal metagenomes from healthy subjects without interventions (Control). This analysis strategy allows to distinguish the effects clearly associated with FMT and separate them from random variations in metagenomic data. The obtained results are presented in Figure 2.

**Figure 2.**
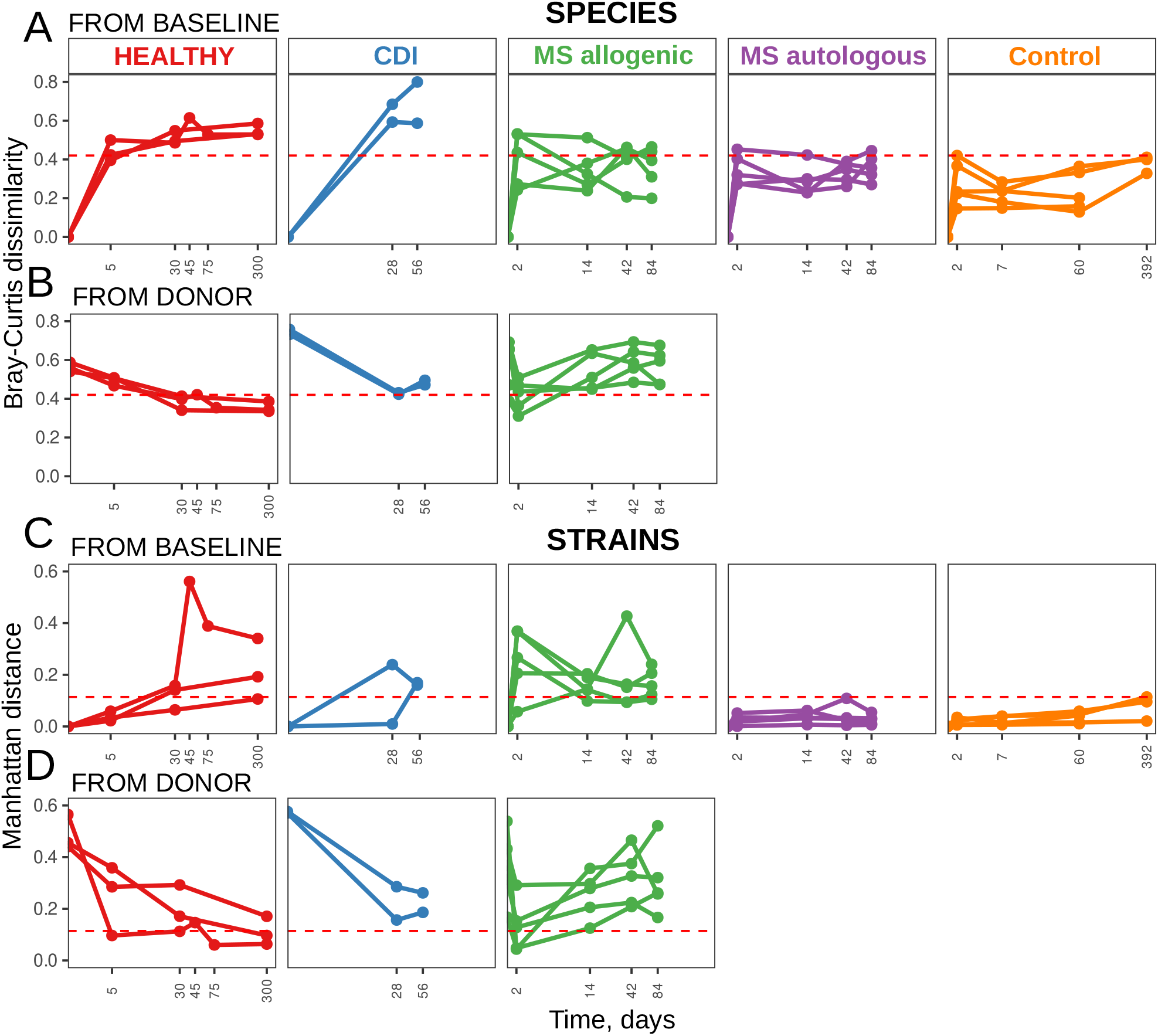
Changes in microbiomes on species (A, B) / strains (C, D) levels due to allogeneic FMT (HEALTHY, CDI and MS Allogeneic groups) above the range of temporal variation observed in autologous FMT (MS autologous group) and healthy controls without interventions (Control). As a comparison metagenome the baseline **(A, C)** or donor **(B, D)** profiles were used. The horizontal red dotted line shows the maximal dissimilarity value between two taxonomic profiles from one person in the Control group.

The species/strains variability over time was statistically significantly higher in the HEALTHY, CDI and MS allogeneic datasets compared to the Control (Wilcoxon rank sum test, *p <* 0.01), while the MS autologous dataset didn’t significantly differs from the Control (Wilcoxon rank sum test, *p >* 0.05). However, the species variability between MS allogeneic and MS autologous sample sets weren’t significant (Wilcoxon rank sum test, *p >* 0.05). The obtained results presented on Figure 2A and 2C. At the same time, the distance based on species and strains levels from the donor sample decreases over time in the HEALTHY and CDI sample sets (see Figure 2B and 2D), while MS allogeneic sample set shows a strong decrease 2 days after transplantation, followed by a gradual increase. Thus, the data obtained indicate a strong effect of the donor microbiota on the gut microbiota recipients profile after FMT in HEALTHY and CDI sample sets, however, in MS allogeneic sample set, this effect is reduced.

#### Sorting metagenomic reads using RECAST algorithm

At this stage, the RECAST metagenomic reads sorting algorithm described earlier was applied. We obtained special reads categories (also pointed as “baskets”) for each time point, which allowed us to dynamically show the changes occurring in the microbiota. Obtained results are presented in Supplementary Figure S2. This figure shows the fraction of classified reads from all sample reads in the corresponding metagenomic samples.

The first “baskets” set (marked **settle** and **not settle**) describes the donor reads that were found/not found in the after-FMT recipient sample. According to obtained results the **settle** reads category represents a smaller fraction of all classified reads. Secondly, pre-FMT recipient reads are searched against post-FMT recipient sample to generate categories **stay** and **gone**. The individual characteristics of the recipients must be pointed. The V1 recipient from HEALTHY sample set showed the almost complete removal of its own metagenomic reads after the second fecal transplant from the same donor, while other recipients from this data set retained a significant part of their own metagenomic reads after transplantation. R02 recipient (CDI dataset) pre-FMT metagenomic sample contained a large number of reads mapped to the human genome, therefore a small number of reads of the bacterial fraction can be classified.

Thirdly, post-FMT sample is separated on four “basket” categories: came **from both** — reads found in both donor and pre-FMT samples, came **from donor** — reads found in donor sample but not found in pre-FMT, came **from baseline** – reads found only in pre-FMT sample and came **itself** – reads not found in donor nor in pre-FMT samples. The results are consistent with the data presented in Figure 2. Recipients from HEALTHY and CDI datasets demonstrated the majority presence of the donor fraction in post-FMT samples, while MS allogeneic recipients showed a conflicting behavior. Only, FAT 008 recipient contains some donor reads fraction in after-FMT samples over time. In other recipients from this dataset this amount is initially small and gradually disappears.

#### Definition of microbial features associated with gut microbiota restructuring

The taxonomic profiling of obtained “baskets” was performed using MetaPhlAn2 [Truong et al., 2015]. As a control (Control data), metagenomic samples of people without interventions were used. The reads were sorted as real-FMT samples, each baseline sample was used as a “donor” sample (see Materials and methods section). This approach will allow to control the effects clearly associated with fecal transplantation.

Based on non-metric multidimensional scaling (NMDS) visualization, the **settle** and **stay** “baskets” form a separate cluster from **gone** and **not settle** categories (see Figure 3A). Analysis of taxonomically classified reads separation into different “baskets” shows a clear difference in microbial composition. The separation of reads from one bacteria to different baskets is a minor event (see Figure 3B). Interestingly, the obtained results are similar between real-FMT and Control data, however settle “basket” of Control data substantially less than real-FMT. Thus, it can be assumed, specific separation rules may be described. To identify these rules, the Random forest algorithm was used. Normalized reads quantities of microbial species distributed between “baskets” were used as a predicted variable (this value was used in the analysis presented in Figure 3B). The classification of the baskets pairs **settle/not settle** and **stay/gone** was performed separately. The taxonomy and relative microbial abundances in donor and pre-FMT metagenomic samples were used as predictive features. Additionally, the average relative abundances of microbes from the HMP 2012 dataset, a donor/recipient subject, time point, and dataset metadata were added to the analysis.

**Figure 3.**
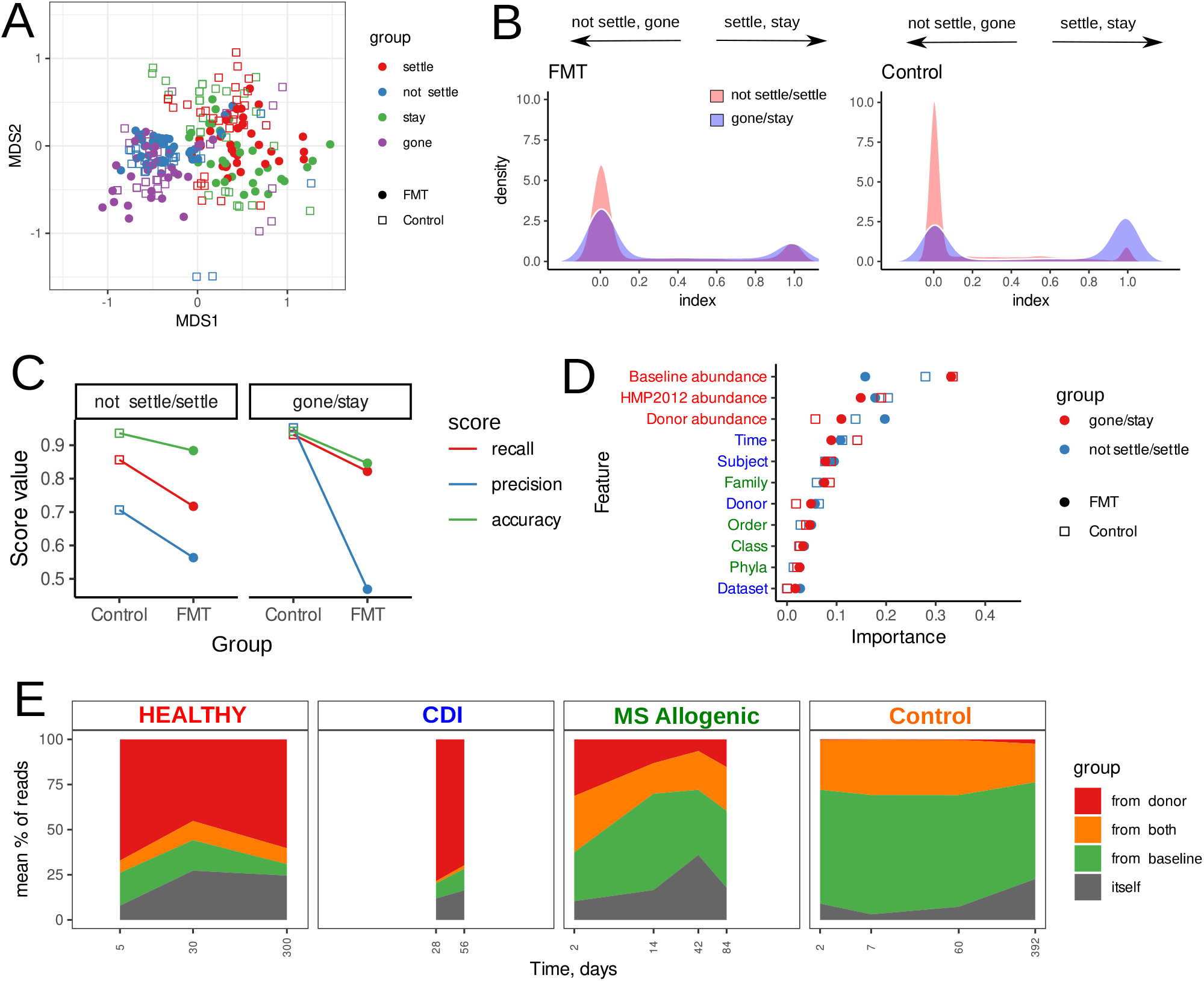
Taxonomic analysis of gut microbiota restructuring during FMT. **(A)** Non-metric multidimensional scaling biplot obtained using species distribution profiles of “baskets” categories and Bray-Curtis dissimilarity. The figure shows the species present in at least 20% of the “baskets” categories. **(B)** Density plot describes microbial distributions in settle-not settle and stay-gone “baskets” categories. Extreme points of the x axis (0 and 1) show the prevailing presence of reads from one bacteria in the corresponding “baskets”. Point 0 corresponds to **not settle** and **gone** categories while the point 1 to **settle** and **stay**. Species whose MetaPhlAn2 markers were covered with less than 100 reads were not included in the analysis. **(C)** Random forest classification quality scores. **(D)** Random forest classification features importance. The groups of features are shown with different colors. The red color corresponds to abundance related features, blue color to metadata features, and green color to taxonomy features. **(E)** Area plots show composition of recipients post-FMT metagenomic samples over time. Percent of reads categories from all recipients were averaged within each group.

The classification quality was lower in the models based on real-FMT sample set in comparison to Control set (see Figure 3C). It may be due to the lack of biological association between samples. However, distribution of features for predicting importance was mostly similar for both “baskets” sets (see Figure 3D). The main contribution to the prediction is made by the all relative abundances variables. However, the influences of the donor variable and donor microbial relative abundances for models based on the Control sample set are reduced. At the same time, the weakest contribution to the prediction is the Dataset variable.

Further, the contribution of each category to the post-FMT metagenomic samples structure was evaluated. For each sample set, the proportion of all categories were averaged. The obtained results are presented in Figure 3E. According to results obtained, HEALTHY and CDI sample sets are mainly formed from the reads coming from the donor, while MS allogeneic mainly from reads coming from their own pre-FMT samples. Also, MS allogeneic is characterized by the presence of a significant **from both** category in comparison with other real-FMT sample sets. It is worth noting that **from both** categories are formed by similar microbial strains, which the sorting algorithm couldn’t clearly distinguish. Also, these can be common parts of the different microbial genomes. This category can be characterized as the donor/recipient common microbiota. Within this group a strains reshaping can occur. Interestingly, the Control sample set analysis shows the high microbiota stability over time. According to this analysis the main part of the microbial diversity comes from the baseline sample, while the complete absence of the microbiota from the “donor”. The **itself** “basket”, apparently, shows random variability, which may be associated with changes in coverage of gut microbiota minor fraction.

#### The restructuring associated with resistance to antibiotics and lantibiotics

The functional capacity of obtained “baskets” has been estimated. The HUMAnN2 pipeline [Franzosa et al., 2018] allowed us to identify 5199 KEGG orthology groups (KO) in all “baskets” categories. Using NMDS visualization, the **settle** and **stay** “baskets” categories form a separate cluster from **not settle** and **gone**(see Figure 4A). To identify functional capacity differences between categories the Songbird method [Morton et al., 2019] was used.

**Figure 4.**
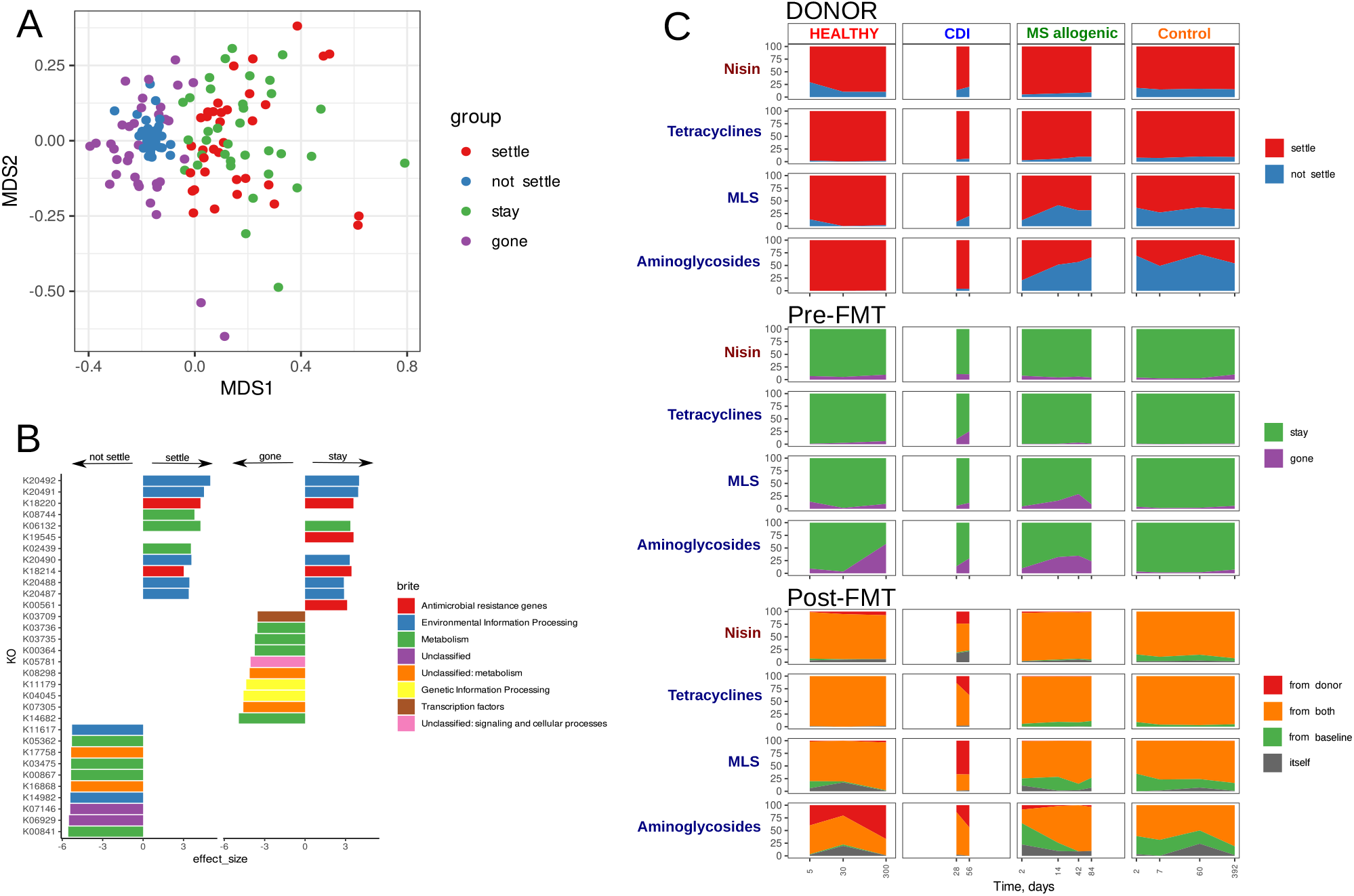
Functional analysis of gut microbiota restructuring during FMT. **(A)** Non-metric multidimensional scaling biplot obtained by using KOs distribution profiles of “baskets” categories and Bray-Curtis dissimilarity. **(B)** The KOs differentially distinguish “baskets” categories. The KOs with negative effect size are associated with **not settle** and **gone**, while **settle** and **stay** KOs have positive effect size. **(C)** Distribution of antibiotic and nisin resistance genes in “basket” categories over time. The related metagenomic sample for all categories is indicated by upper inscriptions. The antimicrobial group of genes is indicated by left side inscriptions. The “basket” categories are indicated by different colors.

Functional differences were determined strictly between dependent “baskets” sets, between **settle/not settle** or **stay/gone** separately. Obtained results are presented in Supplementary Table S4 ans Supplementary Table S5. The analysis showed that the **settle** category was associated with 35 KO while **not settle** category with 150 KO. The top 10 KOs effect size associated with “baskets” shows on Figure 4B. Comparison of **stay** and **gone** categories showed association with 19 and 68 KO respectively. Interestingly, the most part of KO associated with the **settle** and **stay** group are participating in lantibiotic/antibiotic resistance. The top 20 of overrepresented KOs in both groups included K20492 (nisG), K20491 (nisE), K20490 (nisF), K20488 (nisR), K20487 (nisK), K18220 (tetM-tetO), K18214 (tetP A-tet40), K19545 (lnuA C D E), K19299 (aph3-III), and K05593 (aadK). Further, extended analysis of “baskets” antibiotic resistance genes (using MEGARes antibiotic resistance genes database) and nisin (using 5 KOs from KEGG database described above) enrichment in obtained “baskets” was carried out (see Figure 4C).

According to the results obtained, the Control sample set showed a similar effect: the tetracyclines/nisin resistance genes overrepresented in **settle** and **stay** categories with comparison to **not settle** and **gone**. Interestingly, **from both** reads groups of tetracyclines and nisin resistance genes contribute more to the post-FMT in comparison to other “basket” categories. It is worth noting that post-FMT metagenomic samples from HEALTHY and CDI sets receive a new ARGs diversity from donors, while in MS allogeneic groups this almost doesn’t happen.

## Discussion

Fecal microbiota transplantation (FMT) is a great model for studying the ecology of host-associated microbial communities. At the moment, the mechanism of the recipient intestine colonization by the donor microorganisms is a complex problem for microbiology and medicine. Moreover, the question of microbial community restructuring due to FMT is complex and important. How do donor-derived and recipient-derived microbes contribute to microbiota re-assembly after FMT? The most researchers tried to answer this question using comparative analysis of gut microbiota taxonomy of donors and recipients [Shahinas et al., 2012; Hamilton et al., 2013; Seekatz et al., 2014; Weingarden et al., 2015; Mintz et al., 2018; Staley et al., 2019]. The authors noted the shift of recipients’ profiles after FMT towards the donors’ ones. However, these studies didn’t reveal the movement mechanism. Whether this effect is caused by engraftment of donor microbiota or growth burst of the recipient’s own donor-related intestinal microorganisms? It has been important to establish the fact of donor microorganisms colonization of the recipient’s intestine [Li et al., 2016; Lee et al., 2017; Kumar et al., 2017; Smillie et al., 2018]. Moreover, colonization of the healthy volunteers intestines by healthy stool microorganisms of healthy donor has been shown [Goloshchapov et al., 2019]. However, only the behavior of donor strains has been demonstrated. What happens to the recipient’s own microbiota?

Here, we suggest the RECAST algorithm for studying the gut microbiota colonization and restructuring processes. This approach is based on the shotgun metagenomic reads sorting process in accordance with their origin. The classification quality allows to distinguish between species, while distinction between closely related strains is difficult. The sorting was performed using time points of recipient metagenomes, which allowed to study the evolutionary dynamics of donor-derived and recipient-derived microbiota in recipient intestine after FMT. Firstly, donor metagenomic samples are separated into two categories (**settle** and **not settle**) containing reads found and not found in post-FMT recipient samples. Secondly, the second “basket” set (marked **stay** and **gone**) describes the pre-FMT recipient reads that were found/not found within the post-FMT recipient sample. Thirdly, post-FMT recipient sample may contain donor-derived reads (came **from donor** category), recipient-derived reads (came **from baseline** category), and donor-recipient common reads (came **from both** category). We must also mention the **itself** category which contains reads whose origin could not be determined. These may be metagenomic reads from the uncovered microbial diversity of the donor. At the same time, these may be microbes from an uncovered recipient microbial diversity that have received a growth boost due to FMT. It can also be transient microflora. Thus, the RECAST method allows to study FMT-related gut microbiota reshaping, evaluating both the engraftment of the donor microorganisms and the restructuring of the recipient’s microbial profile.

Using the RECAST sorting approach the shotgun metagenomic data from three FMT studies including healthy volunteers [Goloshchapov et al., 2019], patients with clostridial colitis [Lee et al., 2017] and metabolic syndrome [Vrieze et al., 2012] were analyzed. According to taxonomic profiling, the **settle** and **stay** “baskets” are similar and differ from **not settle** and **gone** categories respectively. The intersection between paired baskets (for examples, between **stay** and **gone**) is poor. Thus, the microbial diversity colonization and preservation process can be characterized by the “all-or-nothing” principle according to Smillie et al., 2018. This indicates both the high sorting quality and also specific biological processes causing this distinction.

Using microbial data such as taxonomy, relative abundance in donor/recipient non-sorted metagenomic samples and average relative abundance in HMP2012 data as well as donor/recipient subject, time point and dataset variables, the Random forest classifiers predicting the microbial reads content in “baskets” were constructed. The classification of the baskets pairs **settle/not settle** and **stay/gone** was performed separately. Thus, contribution of described features to the microbial diversity colonization and preservation process was evaluated. According to this analysis, the microbial relative abundance in donor/recipients and HMP2012 metagenomes were the most important features for classification bacteria by “basket” categories. In other words, microbes of **settle** and **stay** categories are associated with the top gut microbial diversity of the human population. Moreover, the Dataset feature makes the least significant contribution for classification bacteria by “basket” categories. Thus, the microbiota of recipients have a common property associated with the process of microbial diversity colonization and preservation regardless of the disease. Similar classifying results in Control and real-FMT sample sets are worth noting. It also indicates the existence of a fundamental rule for the microbial diversity allocation in the human gut.

The results described here are consistent with the previous studies. The donor metagenome-assembled genomes that colonized both recipients were prevalent, and the ones that colonized neither were rare across the participants of the Human Microbiome Project samples [Lee et al., 2017]. Another research stated that engraftment can be predicted largely from the abundance and phylogeny of bacteria in the donor and the pre-FMT patient [Smillie et al., 2018]. Thus, the main rule of colonization can be defined: the top of the donor stool microbiota and likely the top of the human population stool microbiota can colonize the recipient intestine. This rule can be supplemented: the recipient tends to retain top gut microbiota. The bold assumption may be that the top gut microbes form the “core” of the human gut microbiota. In other words, FMT modifies the recipient “core” gut microbiota by the “core” of the donor gut (stool) microbiota. Apparently the “core” microbiota have specific properties which allows microbial “cores” integration and prevent colonization by transient bacteria.

Sorting of post-FMT metagenomic samples have shown donor-derived and recipient-derived microbial diversity distribution and evolution over time. The HEALTHY and CDI sample sets demonstrate a dominance of donor microbiota diversity. However, the recipient’s own microbiota is prevalent in MS allogeneic samples set, while the number of detected common metagenomic reads is increased. Perhaps, donor-derived and recipient-derived microorganisms competed and re-assembled recipient microbiota structure. Also, the host immunity can participate in this process. Thus, the more competent donor/recipient microbes will gain an advantage over the recipient/donor bacteria, which will affect the post-FMT metagenome profile. It could be related to a super-donor effect [Wilson et al., 2019]. Moreover, post-FMT taxonomic profiles can be predicted using taxonomic and abundance data of donor/recipient metagenomic samples [Smillie et al., 2018]. In other words, the re-assembly process also must be related to specific biological rules. In post-FMT samples of Control dataset “donor” microbiota was not found, while the intersection between the “donor” and recipient microbial diversity (described **from both** “basket” categories) was ~ 25%. It is important that the ratio of the categories is relatively stable. Thus, we can conclude that the gut microbiota has a stable unique structure and intersects up to ~ 25% between two people. It can be both overlapping taxonomy and/or common functional features that may be required for the human gut microbiota.

The analysis of functional differences has shown enrichment by nisin, tetracycline, lincosamides, and aminoglycosides resistance genes in **settle** and **stay** baskets. These observations may be associated with previously described colonization resistance phenomena [Bohnhoff and Miller, 1962; Lawley and Walker, 2013; Zmora et al., 2018; Litvak et al., 2019]. According to this hypothesis the definite antibiotics can be produced by gut microbiota itself and form one of the resistance mechanisms against colonization by third-party bacteria. In other words, these genes are part of the recognition system “friend or foe” responsible for getting inside into a “core” microbiota. It is worth noting that *Blautia obeum* is a producer of nisin O which was isolated from human gut microbiota. This research adds to the evidence that lantibiotic production may be an important trait of gut bacteria [Hatziioanou et al., 2017]. Moreover, gut microbiota produces broad spectrum of antimicrobials [Garcia-Gutierrez et al., 2019], which can be included in the development of protection mechanisms against colonization by pathobionts and others random third-party bacteria [Kamada et al., 2013; Kim et al., 2019]. Likewise, tetracyclines antibiotic resistance genes were found within the Hadza hunters-gatherers population in Tanzania, which were not exposed to anthropogenic pressure in comparison to the residents of modern urban areas [Rampelli et al., 2015]. This additionally confirms the ecological role of these genes in the human intestine microbiota. On the other hand, this may be an additional characteristic coupled with other specific features associated with “core” bacteria and colonization resistance. Moreover, apparently it is a common characteristic between donors and recipients (and controls) included in the study. The AR-genes accumulation in “core” microbiota can be caused by systematic exposure to the food-borne antibiotics. It is known that various antibiotics are allowed in many countries’ food additives including nisin [de Arauz et al., 2009].

To summarise, RECAST algorithm offers an improved approach for analysis of FMT experiments metagenomic data which allows researchers to gain novel biological insights. The idea of reads sorting prior to the comparison of metagenomes with the common computational approaches enables to preserve more data by presenting it unchanged from, for example, compared to the metagenomic assembly, which can cause the loss of a significant part of information [Olekhnovich et al., 2018; Brown et al., 2020]. Using the RECAST algorithm, not only FMT data such as [Draper et al., 2018] can be analyzed, but also, for example, the colonization of the organism of children by the microbiota of mothers [Ferretti et al., 2018] or others. This can reveal “hidden” significant biological trends. Further disclosure of the “core” microbiome concept can be used for experimental design development which can be based on a comparison of “core” stool metagenomes between different groups, rather than metagenomic samples. In this case, the variable part of the microbiota can be discarded. This can solve the problem of noise filtering and data dimensionality reduction. Also, this biological insight may supplement existing mathematical techniques. However, this assumption certainly requires additional confirmation. The description of ecological processes underlying the development and existence of host-associated microbial communities can open the way to point-editing microbiota of the human intestine. This can become the basis for the creation of medicines with fundamentally novel mechanisms of action.

## Conclusions

Here, we evolved the previously proposed “core” gut microbiome concept. “Core” metagenome can be defined using relative abundance and taxonomy data. Also, we hypothesize the mutual integration of “cores” microbiota between human individuals. Another “core” microbiota feature is an increased abundance of antibiotic and lantibiotic resistance genes which may be due to both the anthropogenic impact and/or the ecological role of these genes in the human gut microbial community.

## Supplementary materials

**Supplementary Figure S1 Simulation results. (A)** Dependence of bases quality phred score on the position in read for simulated datasets. **(B)** Accuracy on simulation dataset from species with exponential relative abundance distribution. **(C)** Relative abundance of species drawn from exponential distribution. **(D)** Accuracy on simulation dataset from species with uniform relative abundance distribution. **(E)** Accuracy on simulation dataset from strains with different Mash distances.

**Supplementary Figure S2 Distribution of reads categories (“baskets”) in related metagenomic samples (indicated by upper inscriptions).** Lines correspond to recipients, while columns are “basket” categories. The datasets are highlighted in color and **(A,B,C)** symbols.

**Supplementary Table S1** Bacterial genomes used for sorting algorithm testing.

**Supplementary Table S2** Summary data about the donors and recipients across datasets.

**Supplementary Table S3** Metagenomic data preprocessing statistics.

**Supplementary Table S4** KEGG orthology groups are differentially abundant between the settle and not settle “basket” categories.

**Supplementary Table S5** KEGG orthology groups are differentially abundant between the stay and gone“basket” categories.

## Availability of data and materials

The study used data from open sources, which are available at NCBI Sequence Read Archives under the BioProjects accession numbers PRJNA510036, PRJEB12357, PRJNA353655, at European Nucleotide Archive (ENA) database ERP009422 and at https://www.hmpdacc.org. Source code of RECAST algorithm can be found at https://github.com/ivartb/FMT. The analytical scripts and obtained data are available at https://github.com/RCPCM-GCB/RECAST_project.

## List of abbreviations

FMT: fecal microbiota transplantation.
IBD: inflammatory bowel disease.
GRM: genome-resolved metagenomics.
NMDS: Non-metric multidimensional scaling.
CDI: *Clostridioides difficile* infection.
SNV: Single nucleotide variant.
WGS: Whole genome sequencing.
MS: metabolic syndrome.
KEGG: kyoto encyclopedia of genes and genomes.
KO: KEGG orthology.
AR: antibiotic resistance.

## Acknowledgements

We thank Alexander I. Manolov for useful comments, Dmitry E. Fedorov for help in metagenomic data preprocessing and taxonomic annotation, and Andrey E. Samoilov for help in the statistical justification of sorting algorithm.

## Authors’ contributions

**EO**-performed computational experiments using real metagenomic data, interpretation of obtained results, performed data analysis and visualization, contributed to algorithm design, contributed to research idea, and wrote the manuscript. **AI**, **VU**-design and implementation of algorithm, contributed to manuscript preparation. **AI**-performed computational experiments using simulated data, partially wrote the manuscript, contributed to interpretation of the obtained results and data analysis. **EI**-research idea, contributed to interpretation of the obtained results and manuscript preparation.

## Ethics declarations

### Ethics approval and consent to participate

Not Applicable.

### Consent for publication

Not Applicable.

