## Supplementary figures for "Gut microbiota alterations due to fecal transplant"

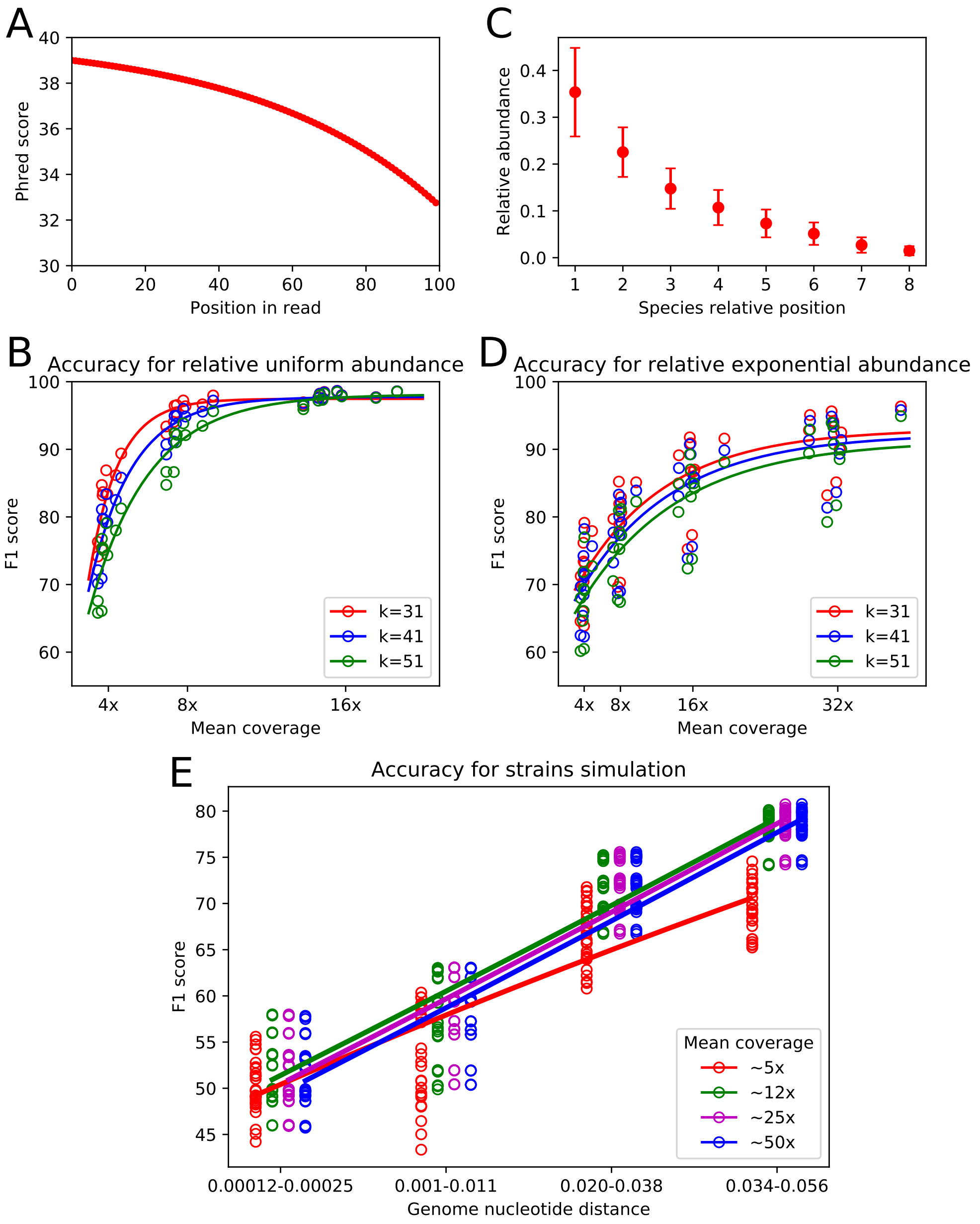


**Figure S1. Simulation results. (A)** Dependence of bases quality phred score on the position in read for simulated datasets. **(B)** Accuracy on simulation dataset from species with exponential relative abundance distribution. **(C)** Relative abundance of species drawn from exponential distribution. **(D)** Accuracy on simulation dataset from species with uniform relative abundance distribution. **(E)** Accuracy on simulation dataset from strains with different Mash distances.

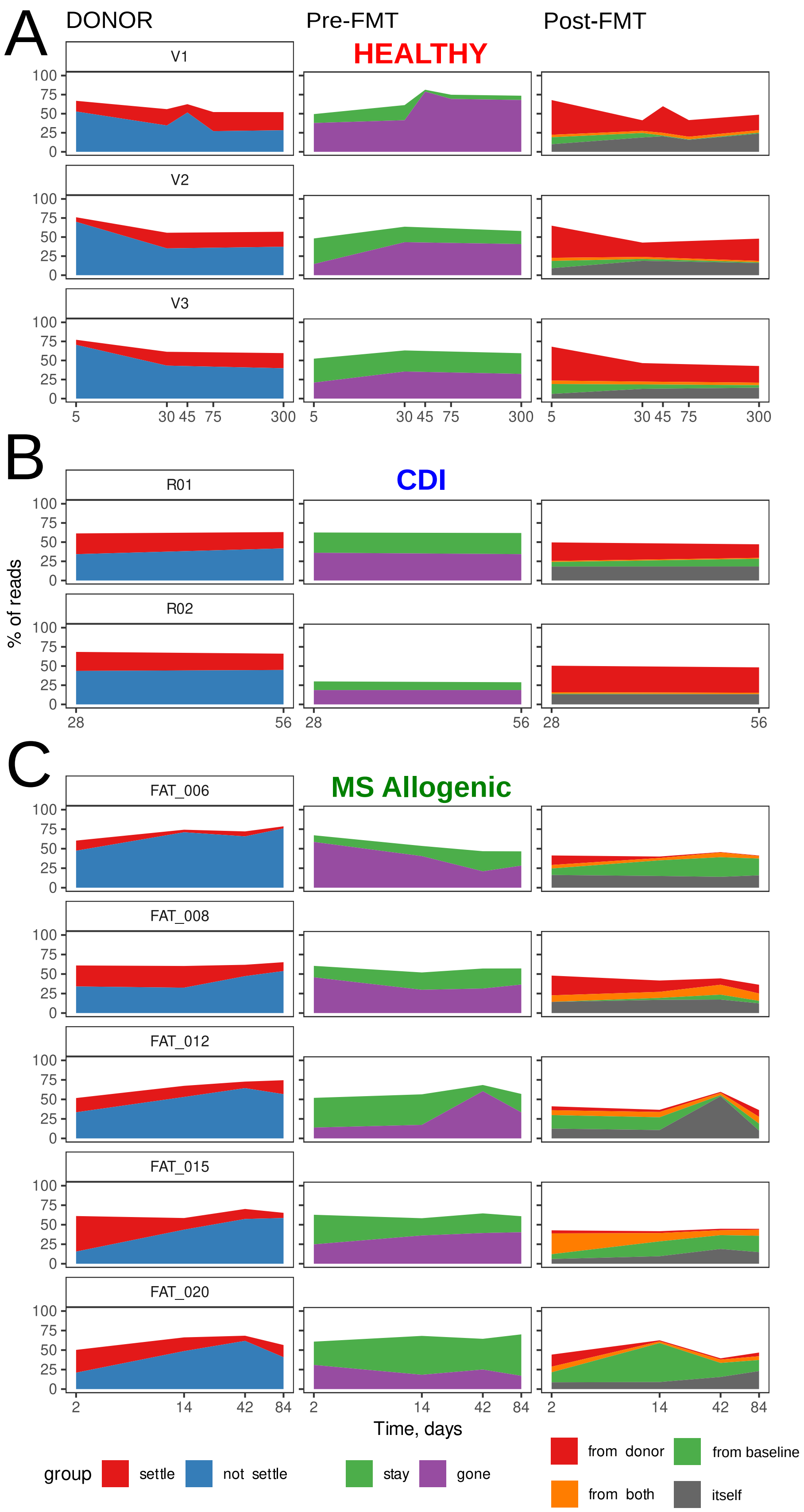


**Figure S2. Distribution of reads categories ("baskets") in related metagenomic samples (indicated by upper inscriptions).** Lines correspond to recipients, while columns are "basket" categories. The datasets are highlighted in color and **(A,B,C)** symbols.
